# Global analysis of Poales diversification – parallel evolution in space and time into open and closed habitats

**DOI:** 10.1101/2023.09.14.557735

**Authors:** Tammy L Elliott, Daniel Spalink, Isabel Larridon, Alexandre Rizzo Zuntini, Marcial Escudero, Jan Hackel, Russell L Barrett, Santiago Martín-Bravo, José Ignacio Márquez-Corro, Carolina Granados Mendoza, Aluoneswi C. Mashau, Katya J Romero-Soler, Daniel A. Zhigila, Berit Gehrke, Caroline Oliveira Andrino, Darren M Crayn, Maria S Vorontsova, Félix Forest, William J Baker, Karen L Wilson, David A Simpson, A Muthama Muasya

## Abstract

- Poales are one of the most species-rich, ecologically and economically important orders of plants and often characterise open habitats, enabled by unique suites of traits. We test the hypotheses that Poales species are assembled into distinct phyloregions, with centres of high phylogenetic diversity and endemism clustered in tropical regions, and that cosmopolitan families show parallel transitions into open and closed habitats at different times.
- We sampled 42% of Poales species and obtained taxonomic and biogeographic data from the World Checklist of Vascular Plants database, which was combined with open/closed habitat data scored by taxonomic experts. A dated supertree of Poales was constructed. We integrated spatial phylogenetics with regionalization analyses, historical biogeography, ancestral state estimations, and models of contingent evolution.
- Diversification in Poales and assembly of open and closed habitats result from dynamic evolutionary processes that vary across lineages, time, space, and traits, most prominently in tropical and southern latitudes. Our results reveal parallel and recurrent patterns of habitat and trait transitions in the species-rich families Poaceae and Cyperaceae, yet other smaller families display unique evolutionary trajectories.
- The Poales have achieved global dominance via parallel evolution in open habitats, with notable, spatially and phylogenetically restricted divergences into strictly closed habitats.

## Introduction

Open and closed habitats are functionally distinct categories of terrestrial environments (Bond, 2019), each with unique ecology and differing in ground layer vegetation strongly constrained by available light (Ratnam *et al*., 2011; Bond, 2022). Open habitats, comprising nearly 60% of land area (Dinerstein *et al*., 2017), include treetops that provide suitable habitats for epiphytes, areas too extreme to support trees (e.g., deserts, high altitudes, tundra, rock walls), and grasslands – regions that could potentially support trees but are dominated by grass-like plants because of disturbances (Strömberg, 2011; Buisson *et al*., 2022).

Grasslands occupy a wide biogeographic range and cover nearly 40% of the terrestrial biosphere, including tropical and subtropical savannas, boreal and temperate prairies, and Eurasian steppes (Bond, 2019; Buisson *et al*., 2022). They provide habitat for a wide diversity of animals and plants, and support the livelihood of over one billion people worldwide (Buisson *et al*., 2022). Until now, biogeographical and evolutionary studies of open habitats have focussed on regional scales (e.g., Solofondranohatra *et al*., 2020) or on the most diverse family occupying these habitats, Poaceae (e.g., Edwards *et al*., 2010; Linder *et al*., 2018; Gallaher *et al*., 2022), although other closely related families can be equally important components of their respective habitats (e.g., Cyperaceae; Barrett, 2013).

The ability of plants to occur in either open or closed habitat has evolved repetitively across the tree of life, in some cases possibly through parallel evolution – the repeated evolution of similar traits in closely-related lineages (Bailey *et al*., 2017; Durán-Castillo *et al*., 2022) – or through divergent evolution involving large evolutionary distances between populations (or species) in genotypic or phenotypic space (Bolnick *et al*., 2018). As the morphological and physiological traits of plants are linked to their habitat preferences, the repetitive evolution of traits suggests parallel adaptive evolution (Bolnick *et al*., 2018). To identify macroevolutionary processes, such as whether evolution is parallel or divergent, it is important to determine the appropriate phylogenetic scope to avoid excluding key lineages that unveil important patterns (Folk *et al*., 2018). Large comparative studies including thousands of species across a single higher-level lineage present opportunity to distinguish which traits might be exceptional at smaller compared to larger phylogenetic scales, as well as incorporate all main clades into the study, including those where the evolution of a trait is typical compared to most of the other lineages (Beaulieu & O’Meara, 2018).

Dominant in open habitats, Poales are among the most species-rich orders of angiosperms, comprising 12 families and 24,302 species (Govaerts, 2022; APG IV, 2016; Fig. 1; Table 1), and including two of the three largest monocot families (Poaceae and Cyperaceae). The 12 Poales families differ in their species richness and distribution, yet plant families exhibiting high species richness tend to be cosmopolitan and varying in life form, such as having both woody and herbaceous species (Ricklefs & Renner, 1994). Although the two largest families are predominately herbaceous, they are cosmopolitan and among the ten most species-rich plant families (Govaerts, 2022). The mid-sized families show latitudinal bias (e.g., Restionaceae in austral temperate regions; Bromeliaceae in the Neotropics) or are restricted to a single continent (e.g., Rapateaceae in South America). The ancestor of Poales is hypothesised to have occupied open and seasonally dry habitats during the Cretaceous (Bouchenak-Khelladi *et al*., 2014) and would have been herbaceous, with typical monocot features including rhizomes, monocarpic tillers, C_3_ photosynthesis, and wind pollination (Dahlgren *et al*., 1985). Several lineages of Poales have entered closed habitats since the Late Cretaceous, but such lineages are predominantly species-poor (e.g., Flagellariaceae), with the exception of Bromeliaceae which has rapidly diversified during the Miocene (Givnish *et al*., 2011, 2014). Conversely, several species-rich Poales lineages have diversified in open habitats, apparently in conjunction with Crassulacean Acid Metabolism (CAM) and C_4_ photosynthetic systems (Bouchenak-Khelladi *et al*., 2014). This generalisation, however, is based on incomplete sampling (below 2% of species; Bouchenak-Khelladi *et al*., 2014), which is likely to obscure underlying patterns and processes, and thus remains to be tested with more comprehensive sampling. From an ancestor occuring in open habitats, multiple lineages of Poales have evolved traits possibly in parallel or divergently to occupy distinct niches.

**Figure 1.**
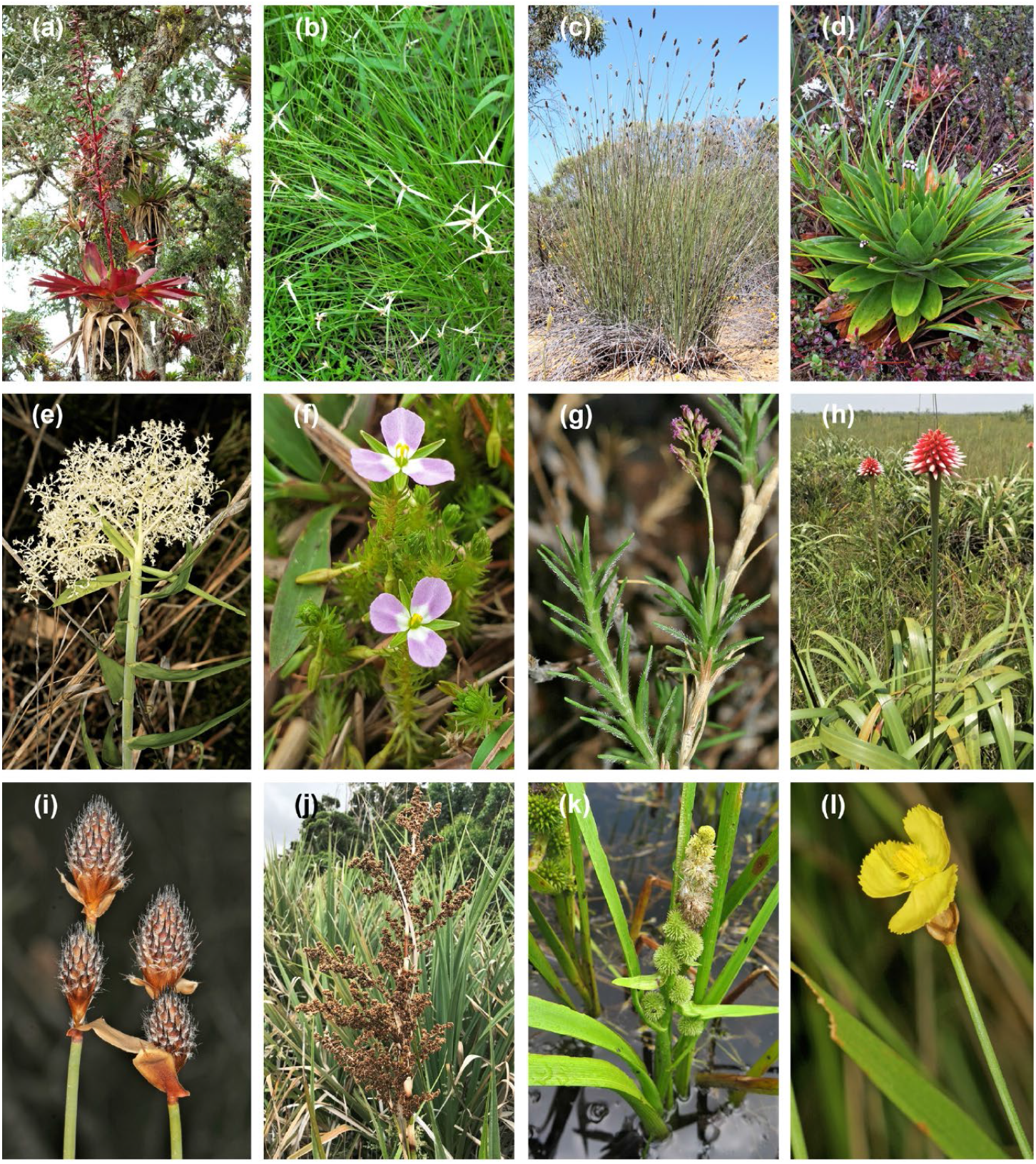
Representatives of twelve Poales families. **(a)** *Tillandsia tovarensis* (Bromeliaceae), epiphytic in cloud forest, Kuelap, N Peru. **(b)** Insect-pollinated *Rhynchospora colorata* (Cyperaceae), in forest gaps, S Ecuador. **(c)** *Ecdeiocolea rigens* (Ecdeiocoleaceae), arid heath, SW Australia. **(d)** *Paepalanthus ensifolius* (Eriocaulaceae), in cloud forest, Podocarpus National Park, S Ecuador. **(e)** *Flagellaria indica* (Flagellariaceae), rocky savannah, NW Australia. **(f)** *Mayaca fluviatilis* (Mayacaceae), wetlands, Singapore. **(g)** *Micraira* sp. Purnululu (Poaceae), a rapid-resurrection species from sandstone pavements in NW Australia. **(h)** *Guacamaya superba* (Rapataceae) in a sedge and grass swamp in E Colombia. **(i)** *Lepidobolus preissii* (Restionaceae) from sandy heath, SW Australia. **(j)** *Prionium serratum* (Thurniaceae) in fynbos, Cape Province, South Africa. **(k)** *Sparganium japonicum* (Typhaceae) from wetlands in E Russia. **(l)** *Xyris complanata* (Xyridaceae) from an ephemeral wetland in savannah, NW Australia. Photos by Russell Barrett; except for **f**, **h**, **j** & **k**, all posted on iNaturalist as CC-BY-NC: **f** by CheongWeei Gan; **h** by Carlos Eduardo; **j** by Linda Hibbin; **k** by Sergei Prokopenko.

**Table 1.**
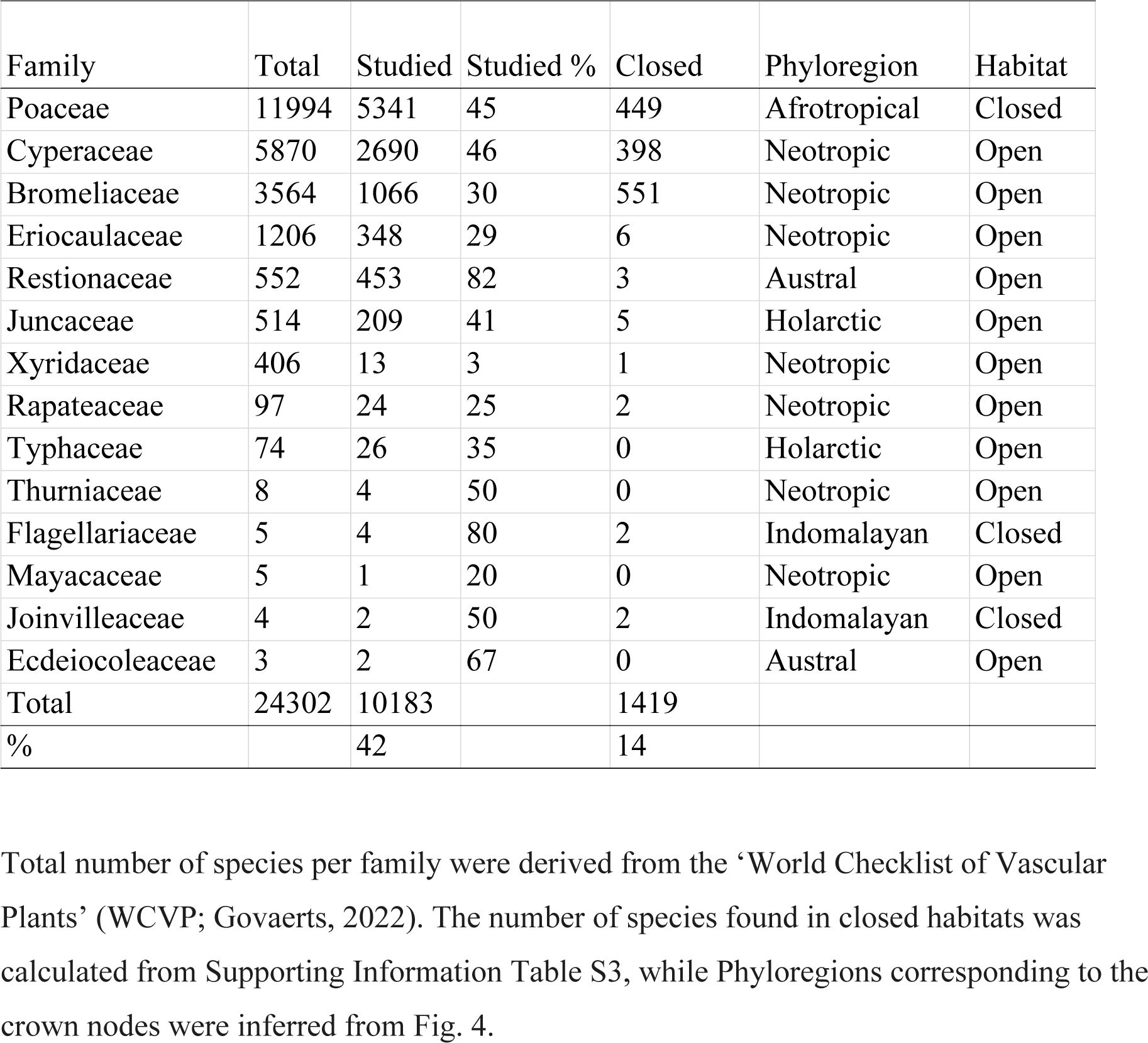
Summary of number of species, habitat and phyloregions corresponding to crown nodes for Poales.

In recent decades, spatial phylogenetic analyses (Mishler *et al*., 2014) have been increasingly used to evaluate the geographic structure of lineage assembly (Holt *et al*., 2013; Thornhill *et al*., 2016; Zhang *et al*., 2022), to link evolutionary processes to the manifestation of biodiversity through time and space (Carter *et al*., 2022; Nitta *et al*., 2022; Sanbonmatsu & Spalink, 2022), and for the development of conservation policies (Sechrest *et al*., 2002; Gonzalez-Orozco *et al*., 2014; Zhang *et al*., 2021). While many spatial phylogenetic metrics exist, emerging methods can identify hotspots of remarkable phylogenetic diversity and distinguish between areas of significant palaeo- and neo-endemism (Mishler *et al*., 2014).

Such analyses provide insight into the spatial patterns of evolutionary processes, integrating phylogenetic information (i.e., branch lengths and species relationships) with the ranges of species and clades. When coupled with functional traits, this inclusion of evolutionary information provides a more complete understanding of macroecological processes (Tucker *et al*., 2017; Spalink *et al*., 2018).

To understand how Poales diversity has accumulated in space and time in the context of open and closed habitats, we focus on the six largest families (Bromeliaceae, Cyperaceae, Eriocaulaceae, Juncaceae, Poaceae, and Restionaceae) to link spatial phylogenetics with models of historical biogeography, phylogenetic regionalization, and ancestral state estimations. We investigate where and when each family diversified across continents, how these processes contribute to phylogenetic regionalization, the spatial and temporal patterns of diversification into open and closed habitats, and whether these patterns demonstrate parallel or divergent evolution. We expect patterns of regionalization to exhibit strong partitioning between tropical and temperate zones, reflecting a tropical origin and select temperate diversification of clades. In addition, we predict that evolutionary transitions between open and closed habitats have occurred frequently, rapidly, and in parallel in Poales, as three of the largest families (Poaceae, Cyperaceae, and Bromeliaceae) have diversified in both habitats. We also expect that evolutionary transitions to, and diversification within, open habitats has occurred in synchrony around the world and across lineages, with the rapid accumulation of species beginning in the Eocene as open habitats became more widespread (Strömberg, 2011). The manifestation of these processes on spatial patterns of diversity should result in significantly high phylogenetic diversity and endemism across the tropics, where most Poalean families have a hypothesised origin, and significantly low endemism in northern temperate habitats. Across Poales, we hypothesise that centres of neo-endemism occur on young islands and mountainous regions, whereas palaeo-endemism should occur in areas with high bioclimatic stability since the Eocene. Finally, we expect family-level spatial phylogenetic patterns to reflect the historical biogeographic context of each clade, with partitioning of neo- and palaeo-endemism dependent on when and where each family diversified.

## Materials and Methods

### Phylogenomic backbone reconstruction

We produced a family-level phylogenomic backbone using nuclear data from 353 loci (Angiosperms353; Johnson *et al.,* 2019). The sampling for the backbone aimed towards 50% of the currently accepted genera and involved new data produced and samples mined from public repositories. The genomic data production was conducted following Baker *et al*. (2022), with DNA extractions (mostly from herbarium materials) using CTAB (Doyle & Doyle, 1987). We used the NEBNext Ultra II DNA Library Prep kit (New England Biolabs) for standard pair-ended library preparation and libraries were hybridised with myBaits Angiosperms353 v1 probe kit (Arbor Biosciences).

The sequence recovery from raw data (target enrichment and mined reads) started with reads being trimmed for short and/or low-quality sequences using Trimmomatic (Bolger *et al*., 2014) and then assembled with a *de novo* approach implemented in HybPiper v.1.3.1 (Johnson e*t al.*, 2016). In HybPiper, trimmed reads were initially binned into genes using BLASTN, which were assembled into scaffolds using SPADES (Bankevich *et al*., 2012), and the coding regions later extracted with Exonerate (Slater & Birney, 2005). For assembled datasets (i.e., whole genomes and transcriptomes) sequence recovery followed Baker *et al*. (2022). Only coding regions were used for the analysis.

We inferred the phylogenomic backbone using a multi-species coalescent framework (MSC). For that, we computed individual gene trees, each computed as follows. The sequences were aligned in MAFFT (Katoh & Standley, 2013) in einsi-mode, with gappy sites (> 90% missing data) removed using Phyutility (Smith & Dunn, 2008). The gene tree was inferred using IQ-TREE 2 (Minh *et al*., 2020), with support assessed via UltraFast bootstrap (UFBS; Hoang *et al*., 2018). Once all gene trees were computed, we used TreeShrink (Mai & Mirarab, 2018) to identify outliers that significantly increased the tree space. For the gene trees with outliers, a new iteration of sequence alignment, trimming and tree inference was performed as described above for the remaining sequences. Once the second iteration of gene tree inference was completed, all gene trees were trimmed for poorly supported branches (UFBS < 30%) and used as input for the MSC analysis in ASTRAL-III (Zhang *et al.,* 2018). In order to obtain a species tree with branch lengths proportional to the genetic distance, we first ranked the genes according to the congruence of their resulting trees to the species tree using SortaDate (Smith *et al.,* 2018) and then concatenated the alignments of the 25 most congruent genes. Using the MSC species tree as topological constraint and this concatenated alignment, a new phylogram was inferred in IQ-Tree 2. For more details on library preparation and data analyses, please refer to Baker *et al*. (2022).

### Reconstruction of species-level phylogeny

We inferred separate trees for five groups of families identified with the backbone phylogeny: 1) Bromeliaceae + Typhaceae; 2) Rapateaceae + Thurniaceae + Juncaceae + Cyperaceae + Mayacaceae (the cyperid clade); 3) Xyridaceae + Eriocaulaceae (the xyrid clade); 4) Restionaceae; and 5) Joinvilleaceae + Flagellariaceae + Ecdeiocoleaceae + Poaceae. Available sequences for different marker sets (Table S1) were retrieved from GenBank (Benson *et al*., 2010). In general, we downloaded as many sequences as possible per marker while disregarding those corresponding to hybrids and not determined to species level, with the exception of the cyperid clade where we preferentially used sequences included in the phylogenetic reconstruction of Elliott *et al*. (2022). Reconciliation of the GenBank names for sequences with those in the ‘Plants of the World Online’ (POWO, 2022) database was done using the R package *TAXIZE* (Chamberlain & Szocs, 2013). We selected one sequence per marker per taxa, while considering synonymy. Sequence matrices for the five clades were aligned using MAFFT v.7.453 (Katoh & Standley, 2013), edited manually in AliView v.1.26 (Larsson, 2014) and concatenated. Trees were then inferred under a GTRCAT model in RAxML v.8.2.12 (Stamatakis, 2014), using rapid bootstrapping with 100 replicates followed by a thorough maximum likelihood search on the Czech National Grid Infrastructure. The preliminary phylogenetic trees were then manually checked for obviously spurious species placements, as well as nomenclatural abnormalities. At this point, we verified that species nomenclature aligned with the ‘World Checklist of Vascular Plants’ (WCVP; Govaerts, 2022). Corrected matrices were then realigned with outgroup taxa (Table S1) and manually edited. A second set of phylogenetic constructions were then conducted using constraint trees derived from the backbone tree.

The backbone and clade trees were time-calibrated using penalized likelihood as implemented in treePL, with the smoothing value identified through cross-validation (Smith & O’Meara, 2012). We used secondary calibration points for the backbone tree, setting fixed ages for the family crown nodes (Table S2) obtained from the plastome tree of Givnish *et al*. (2018). Separate analyses were conducted for Poaceae and Cyperaceae, incorporating additional fossil and secondary priors within the clades using best available information (Spalink *et al*., 2016; Gallaher *et al*., 2022; see Table S2). For the remaining clade trees, the root ages were set to one. All subtrees were then grafted into the backbone tree after scaling their age to that of the corresponding nodes in the dated backbone tree.

### Species distributions and habitats

We used the WCVP dataset to create a species presence/absence matrix in each botanical country (Level 3) as specified by the International Working Group on Taxonomic Databases for Plant Sciences (TDWG; Brummitt *et al*., 2001). Species not present in the phylogeny were removed from the matrix prior to downstream analyses. Occurrence in open, closed or both habitats was scored from the literature, especially regional floras, in addition to taxonomic expertise. Habitats noted as ‘forests’ in floristic treatments, interpreted as closed canopy at least during the active growing season, were scored as ‘closed’. We scored species occurring in open and closed habitats as ‘both’. For Bromeliaceae, species occurring on sun-exposed rock-walls or treetops were scored as ‘open’. A full list of taxa included in the study and their habitat is available in Table S3.

### Regionalization

To determine phylogenetic patterns of regionalization in Poales, we first calculated phylogenetic beta diversity across all botanical countries. We then used three metrics to identify the optimal number of phyloregions: K-means and silhouette scores (Hartigan & Wong, 1979; Rousseeuw, 1987), the elbow method (Salvador & Chan, 2004; Zhao *et al*., 2011), and the gap statistic (Tibshirani *et al*., 2001). For each analysis, we set the maximum number of phyloregions to 20, allowing for the possibility of high levels of regionalization. The elbow and K-means approach each identified 13 phyloregions as optimal, whereas the gap statistic identified 20 phyloregions. For simplicity, we used 13 phyloregions for downstream comparative analyses, with regions partitioned to the optimal number identified above using non-metric multidimensional scaling (NMDS) and hierarchical dendrogram clustering based on a beta-diversity similarity matrix of botanical countries. To determine nestedness of the 13 regions, we completed a final analysis that limited the number of phyloregions to three. Regionalization analyses were conducted using the R packages *phyloregion* v.1.0.6 (Daru *et al*., 2017, 2020a, 2020b), *cluster* v.2.1.3 (Maechler *et al*., 2022) and *stats* v.4.1.2 (R Core Team, 2021). To summarise the spatial and temporal patterns of lineage assembly, we used the R package *phytools* v.1.2.0 (Revell, 2012) to construct lineage through time plots for the open and closed species of each family in each phyloregion.

### Spatial phylogenetics

We used the R packages *picante* v.1.8.2 (Kembel *et al*., 2010) and *canaper* v.0.0.2 (Nitta & Iwasaki, 2021) to calculate phylogenetic diversity (PD; Faith, 1992) and phylogenetic endemism (PE; Rosauer *et al*., 2009), as well as to perform categorical analysis of neo- and palaeo-endemism (CANAPE; Mishler *et al*., 2014) across botanical countries. To calculate the standardised effect size of PD and PE, we used two different sequential randomization algorithms – swap and curveball. The swap algorithm shuffles species presence/absence across the community matrix, while maintaining species ranges sizes and species richness within communities (Gotelli & Entsminger, 2003). The curveball algorithm identifies the species that occur in only one of two randomly selected communities, then distributes those species across communities, while maintaining species richness in communities (Strona *et al*., 2014). For both randomizations, we performed 1,000 replications of 100,000 iterations each. These were followed by two-tailed tests to determine if PD and PE were significantly over- dispersed or clustered. CANAPE methods follow those outlined in Mishler *et al*. (2014) and Nitta & Iwasaki (2021). For PD, PE and CANAPE, analyses were conducted for all Poales combined, as well as separately for the six largest families (Bromeliaceae, Cyperaceae, Eriocaulaceae, Juncaceae, Poaceae and Restionaceae). In addition, PD was calculated for both open and closed habitat species.

### Ancestral state estimations

The ancestral area estimation provides biogeographical context for lineage shifts between open and closed habitats. Our geographical unit of comparison is the Poales phyloregions identified above, but simplified from 13 regions to 7 to reduce the state-space explored by the model (see inset map in Fig. 4). We *a priori* selected the dispersal-extinction-cladogenesis (DEC) model of ancestral estimation as the model with the most appropriate parameters for our study system (Ree *et al.,* 2005; Ree & Smith, 2008; see Notes S1 for justification), which we implemented with stochastic mapping using the R package *BioGeoBEARS* v1.1.2 (Matzke 2013, 2014). For comparison, we also ran the full suite of DEC, DIVA-like, and BAYAREA- like models available in *BioGeoBEARS*, with and without the additional J parameter. Stochastic maps were summarised to estimate the frequency of dispersals between bioregions.

To estimate ancestral states of open/closed habitat and the transition rates between them, we used Generalised Hidden Markov models, as implemented in the function corHMM in the R package *corHMM* v.2.8 (Boyko & Beaulieu, 2021). We ran “symmetric rates” and “all rates differ” Markov models separately with one transition regime and with two transition regimes associated with two hidden states fitted during the analyses. Akaike information criterion (AIC) was used to select the best fitting model, and ancestral states in the phylogeny were inferred with maximum likelihood and stochastic mapping. We then linked the output of the best-fitting *corHMM* models to the DEC analysis to summarise spatial patterns of open/closed habitat transitions, identifying the nodes at which character states shifted, and the geographical areas that those ancestors were inferred to have occupied based on the DEC analysis.

## Results

We inferred a phylogenetic supertree encompassing 10,100 species of Poales (41.6%; Table 1). All families and 88% of genera in the order were represented, but the percentage of species sampled varied from highest in Restionaceae (82%, 453 species), medium for the largest families Poaceae (45%, 5341 species) and Cyperaceae (46%, 2690 species), to lowest in Xyridaceae (3%, 13 species). The majority of species studied occur in open habitats (vs. 14% in closed). Open/closed habitat scoring was carried out for all species, but with some gaps where information was not available (Table S3; > 80% of taxa scored per family).

### Regionalization and lineages through time

Phylogenetic regionalization identified 13 regions (Fig. 2a), which were nested into three major zones (SI Fig. S1): 1) **Temperate** (1: temperate North America + Russian far east; 2: northern Africa [excluding Algeria] + Arabian peninsula [excluding Yemen] + southwestern Asia; 4: Patagonia + Antarctica + western and southern Australia + New Zealand; 5: central and eastern North America; 6: Europe + mediterranean Eurasia + northern Africa [Algeria]; 8: northeastern Asia; 12: central and eastern China; 13: north central Pacific), 2) **Neotropics** (3: Central + South America [excluding central and eastern Brazil]; 11: central and eastern Brazil), and 3) **Palaeotropics** (7: Sub-Saharan Africa [excluding central and western Africa; Madagascar + Mozambique] + southeastern Arabian Peninsula [Yemen] + Mauritania; 9: Indomalayan, comprising India + southeastern Asia + northeastern Australia + Madagascar + Mozambique; 10: central and western tropical Africa [excluding Mauritania]). A NMDS plot shows these 13 regions separated latitudinally, falling into three groups, where northern and southern hemisphere temperate regions form a single cluster separated from the Holotropics, and the latter separates into Neotropics and Palaeotropics (Figs. 2a, S1).

**Figure 2.**
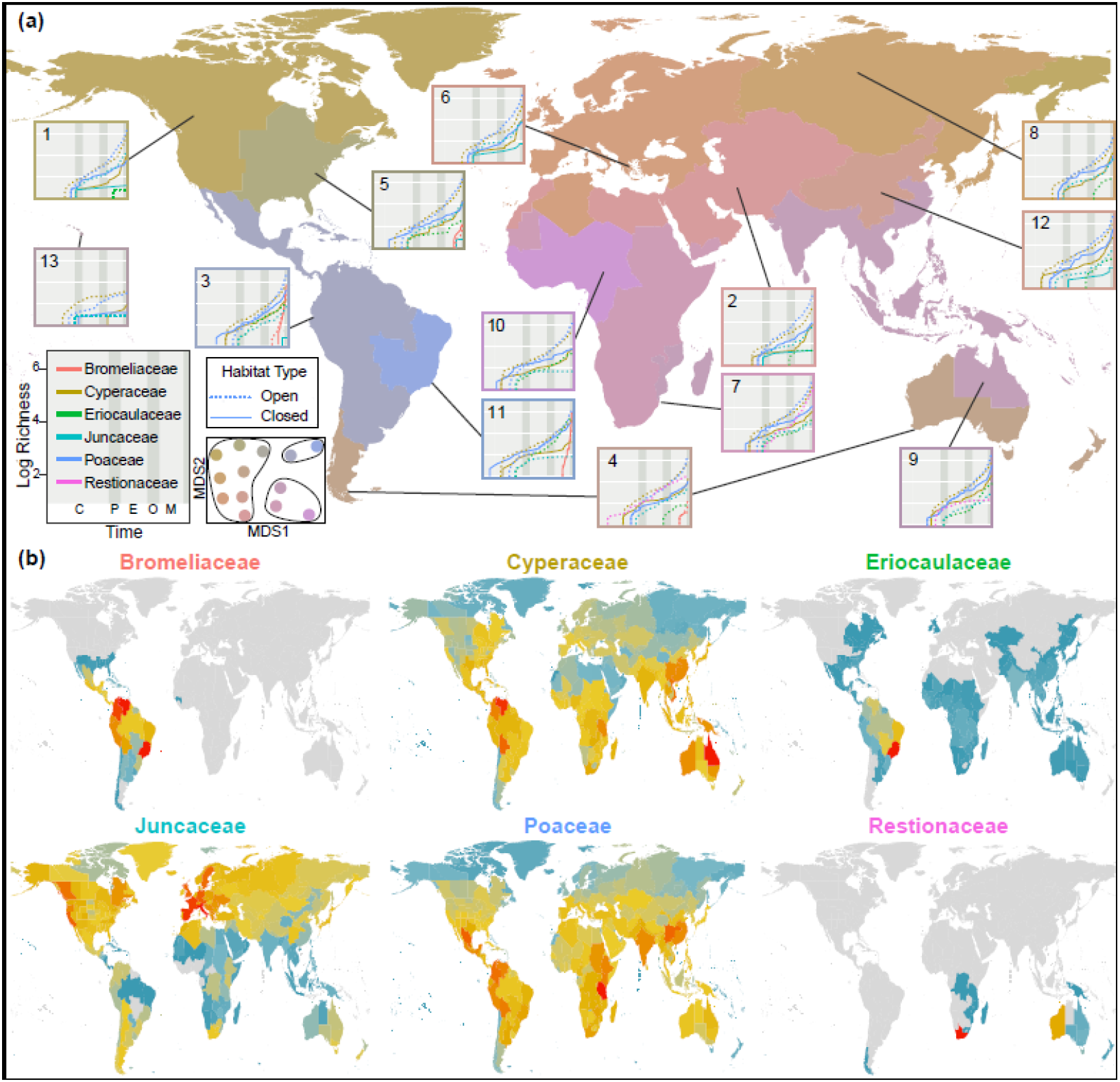
**(a)** Phylogenetic regionalization of the six largest families of Poales. Graphs represent the logarithmic richness of the six families through time within each of the 13 phyloregions identified using the elbow and K-means approach, where dotted lines represent open habitat lineages and solid lines indicate closed habitat lineages. A geological timeline is located in the inset in the far left-hand corner, with the following abbreviations: Cretaceous (C: 66–56 Mya); Paleocene (P: 66–56 Mya); Eocene (E: 56–33.9 Mya); Oligocene (O: 33.9– 23 Mya) and Miocene (M: 23–5.3 Mya). Inset to the right of the geological timeline shows nestedness of the 13 regions based on non-metric multidimensional scaling (NMDS), where each colour dot represent a phyloregion and the three clusters each with similar colour dots represent the three major botanical kingdoms (Temperate, Neotropics, Palaeotropics). **(b)** Phylogenetic diversity (PD) of the six most speciose families of Poales, with red and dark blue indicating botanical countries with relatively high and low PD values, respectively. Botanical regions indicated by grey in **(b)** lack species for the respective family.

Major families differ in their stem age and lineage accumulation over time among the 13 regions (Fig. 2a). When considering the six largest families, either the Poaceae or Cyperaceae are the oldest in all regions, except for the Restionaceae which are oldest in temperate Australia. Bromeliaceae is the youngest family, whereas the small families (Eriocaulaceae, Juncaceae) are younger than Poaceae and Cyperaceae in regions where they occur. Poaceae and Cyperaceae occur in all regions, Juncaceae is nearly cosmopolitan but is poorly represented in tropical regions, while the other nine families are restricted to one or few regions. Considering the six largest families, three are represented in Asia and northern Europe (Poaceae, Cyperaceae, Juncaceae); five occur in North, Central, and South America (Poaceae, Cyperaceae, Bromeliaceae, Eriocaulaceae, Juncaceae); five are found in the Austral, tropical and subtropical Africa regions (Poaceae, Cyperaceae, Eriocaulaceae, Juncaceae, Restionaceae; Fig. 2a,b).

In all tropical regions, Poales – and specifically Poaceae – diversified first in closed habitats before lineages began accumulating in open habitats (Fig. 2a). By contrast, in northern temperate regions open habitat lineages - specifically, Cyperaceae – diversified first (Fig. 2a). The exception to this latter pattern is in the eastern Nearctic, where closed habitat Poaceae were the first to diversify. However, this exception is likely driven by the inclusion of some subtropical Caribbean islands and southern Florida in this bioregion. The initial diversification of Poaceae in most temperate regions included both open and closed habitat lineages. Likewise, but in tropical regions only, the initial diversification of open and closed habitat Cyperaceae was concurrent. Open habitat Poaceae displayed the greatest accumulation of lineages (especially during the Oligocene and Miocene) followed by open habitat Cyperaceae in all but the north central Pacific phyloregion, where lineage diversification was more gradual and where there was a big lapse in time until Cyperaceae diversified in closed habitats (Fig. 2a). The most rapid accumulation of lineages occurred in the closed habitat Bromeliaceae. In all bioregions, open habitat species richness is higher than closed habitat species richness in all families except Bromeliaceae.

### Spatial Patterns of Diversity

The species in our phylogeny were generally representative of the cosmopolitan distribution of Poales, except for the underrepresentation of tropical areas in South America, India and parts of China (SI Fig. S2). Across Poales and in all families except Bromeliaceae, phylogenetic diversity is substantially higher in open habitats than closed habitats in all bioregions (Figs. 3a, SI Fig. S3), with the highest values attributed to open habitat species in southern Africa, western Australia, and northern South America. High open habitat PD is driven by Cyperaceae, Poaceae, and Restionaceae in southern Africa; Cyperaceae, Poaceae, and Restionaceae in western Australia; and by Bromeliaceae, Cyperaceae, Eriocaulaceae, and Poaceae in northern South America (Figs. 2b, 3; SI Fig. S3). Although closed habitat PD is lower overall than open habitat PD, northern South America is a high centre of diversity of these species (Fig. 3b), with high representation of Bromeliaceae, Cyperaceae, and Poaceae. Despite the widespread availability of forested habitats throughout the Holarctic, these regions have low PD of closed habitat Poales. Those species that do occur in closed habitats in the Holarctic represent only Cyperaceae, Juncaceae, and Poaceae (SI Fig. S3). Apart from Juncaceae, closed habitat PD is higher in all major families south of the Tropic of Cancer (SI Fig. S3).

**Figure 3.**
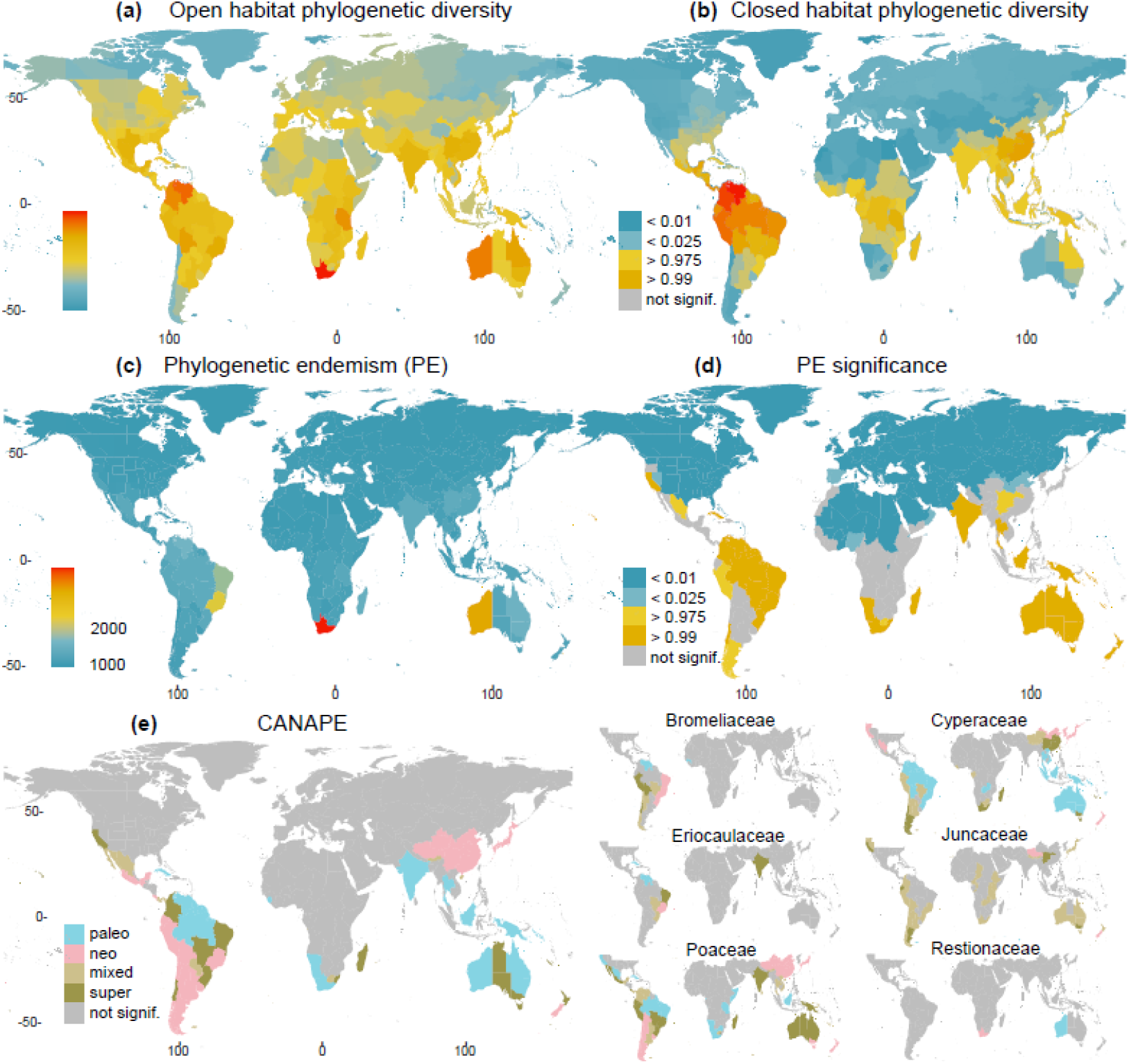
Biodiversity patterns in Poales represented by Faith’s phylogenetic diversity (PD) for open habitat species **(a)** PD for closed habitat species; **(c)** Rosauer’s phylogenetic endemism (PE) and **(d)** PE significance; and **(e)** centres of palaeo- and neo-endemism determined using CANAPE for all Poales (left) and the six most speciose families (right). High PE significance indicates that the region has an overrepresentation of short, rare branches. Low PE significance indicates that short, rare branches are underrepresented.

By far, southern Africa displays the highest PE in Poales (Fig. 3c), driven by high endemism of Cyperaceae, Poaceae, and Restionaceae (SI Fig. S4). PE is also high in Western Australia, again because of restricted lineages from the same three families, and southeastern Brazil, largely due to Bromeliaceae and Eriocaulaceae (Figs. 3c, SI Fig. S4). The Tropic of Cancer clearly demarcates centres of significant PE, with nearly all Holarctic regions exhibiting significantly low PE and many botanical countries in the tropics and subtropics exhibiting significantly high PE. California is the only north temperate zone with significantly high PE, driven by Poaceae and Juncaceae (Figs. 3c, SI Fig. S4). Overall, family level patterns of significant PE match those of Poales, with low PE in the North and high PE in the South, except for Typhaceae which tends to have low PE in most of its range.

Categorical analysis of endemism (CANAPE; Fig. 3e) shows a general contrasting pattern for Poales between Holarctic (mostly non-significant) and Holotropic + Austral (significant) regions. Palaeo-endemism in Poales was supported in northern South America, southern Africa and Australia – a pattern also shared with Poaceae (including Tanzania, California) and Cyperaceae, whereas palaeo-endemism is only evident in Australia for the Cyperaceae and Restionaceae. Neo-endemism is observed in western South America (Andean mountains) in the Poales and Poaceae, but in the eastern South American region (Brazil) for Bromeliaceae and Eriocaulaceae; in most of East Asia for the order and families Poaceae and Cyperaceae; and in southern Africa for the Restionaceae. A mixture of both palaeo- and neo-endemism (i.e., supra-endemism) is observed in Australia and South America for both the Poales and Poaceae. Though areas of endemism in Poaceae and Cyperaceae often co-occur, frequently centres of palaeo- or neo-endemism in one family are centres of mixed/super endemism in the other (e.g., Andean Mountains, Australia, southeast Brazil, eastern China, southern Africa).

### Historical Biogeography and Ancestral Habitat Estimations

The BioGeoBEARS model with the highest AICw was the BayAreaLIKE +J model, followed by DEC without the additional J parameter (Table S4). Given that the BayArea model does not parameterize vicariant speciation but instead allows widespread sympatric speciation, we do not consider it a reasonable model for the global analysis of a clade that spans 120 million years of evolution. Indeed, the BayAreaLIKE model places the Poales in a nearly cosmopolitan range for the first 40 million years of its evolution, which is neither supported by fossil data nor biologically plausible. Thus, subsequent analyses are based on the DEC model. Poales are inferred to have originated in the Neotropics in Western Gondwana (current South America) (Fig. 4a). The crown nodes of most families occur in the Cretaceous (e.g., Cyperaceae, Eriocaulaceae, Juncaceae, Poaceae, and Restionaceae) to the Neogene (e.g., Bromeliaceae) (Fig. 4a). The family crown nodes are distributed in all the major zones (Fig. 4a; Table 1): Bromeliaceae, Cyperaceae, Eriocaulaceae, Rapataceae, and Xyridaceae are assigned a Neotropic origin, while Flagellariaceae and Poaceae are placed in the Palaeotropics. The Holarctic region is ancestral for the Juncaceae and Typhaceae, whereas the Austral region is ancestral for the Restionaceae. Below family rank, Schoeneae is shown to have originated in the Austral region, *Carex* and Pooideae in Eurasia, Cypereae and the PACMAD clade in Sub-Saharan Africa, Abildgaardieae in Indomalayan region (including NE Australia, Mozambique, and Madagascar) and Bambusoideae in the Neotropics (Fig. 4a). Of these clades, Schoeneae is the oldest originating c. 70 Mya, while *Carex* is the youngest with an origin dated at c. 34 Mya.

Poaceae and Cyperaceae experienced a several million-year lag before co-occurring in any bioregion (Figs. 2, 4). This pattern of concurrent but spatially separated diversification is repeated for main lineages within these families as well. In the Paleocene, the grass clades Bambusoideae, PACMAD, and Pooideae originate allopatrically in the Neotropics, Afrotropics, and Eurasia, respectively. In the Eocene, the sedge clades Abildgaardieae, Cypereae, *Carex* and Schoeneae originate in the Indomalayan tropics, Afrotropics, Palearctic, and Austral region, respectively.

Dispersal across all bioregions has occurred many times throughout the evolution of Poales, most frequently between the Neartic and Neotropics and between the Palearctic and Paleotropics (Fig. 4b). Eurasia is a strong one-way source to North America, tropical Africa and tropical Asia, while equatorial Africa is a notable recipient of lineages from Sub-Saharan Africa (Fig. 4a,b). A relatively high number of transitions between open and closed habitats have occurred in the Neotropics and Eurasia, while numbers are low for equatorial Africa and the Austral region (Fig. 4a,c). Transitions from open to closed are higher than the opposite for the Neoarctic and Sub-Saharan Africa, whereas closed to open transitions are higher for the Indomalayan region (Fig. 4a,c).

The asymmetrical (“all-rates-different”) with two rate regimes is the best fit corHMM model of evolution for open/closed habitats (AIC = 8814.2; Fig. 5; Table S5). An open habitat (Fig. 5) is inferred at the root of Poales and most families, with a slow rate shift to closed habitat (0.000000001 transitions per million years [Mya] in R2 [Regime rate 2]).

**Figure 4.**
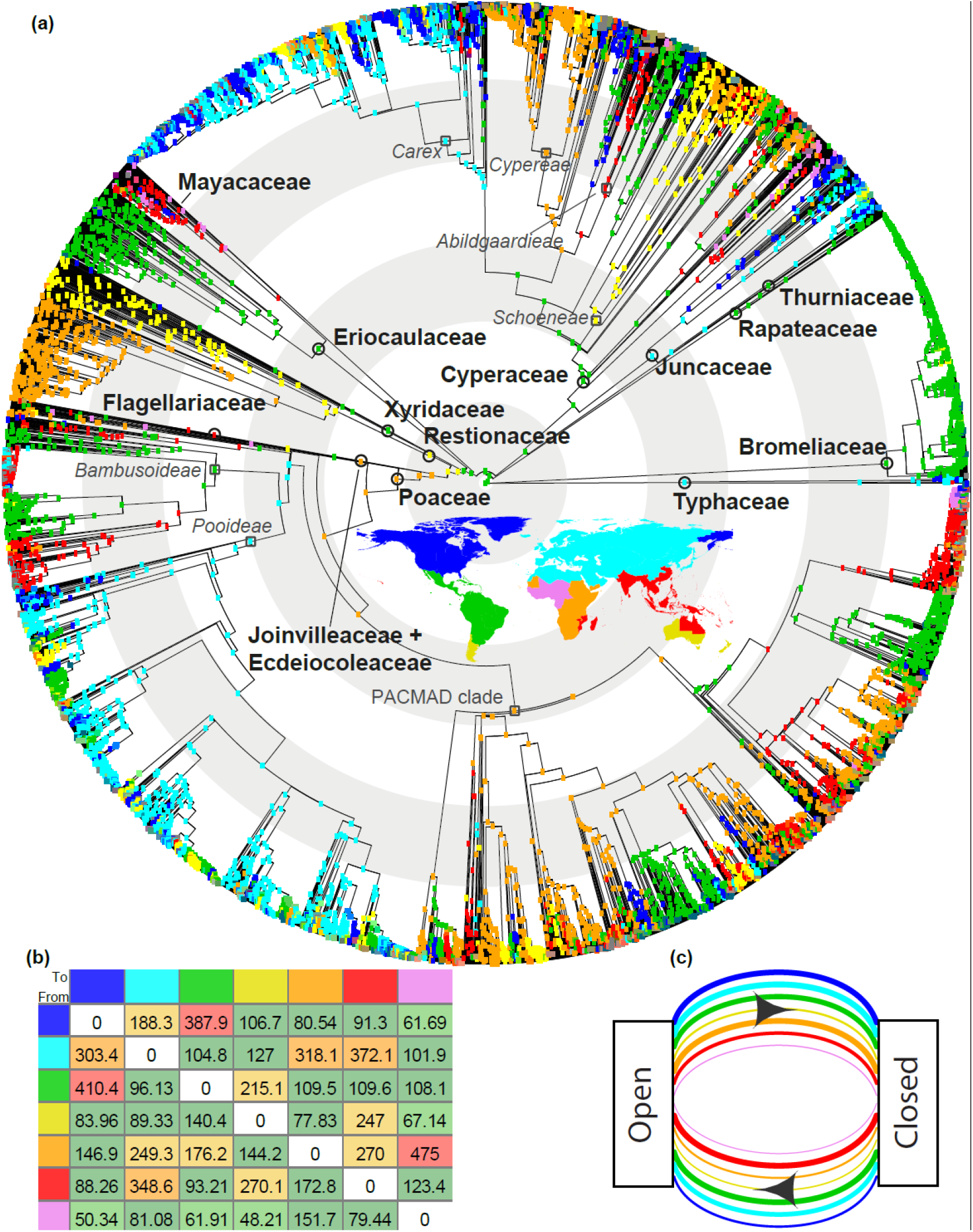
Ancestral area reconstruction within Poales based on seven regions, obtained using the DEC model in BioGeoBEARS **(a).** A global map showing colours corresponding to the seven defined areas for the BioGeoBEARS analysis is in the centre of the phylogenetic reconstruction. The crown nodes of the families within Poales are shown with black circles, whereas dark grey squares are used to depict lineages important for the study’s interpretations. Concentric light grey/white rings underlying the phylogeny indicate time slots of 20 million years intervals. Note that Joinvilleaceae and Ecdeiocoleaceae are depicted together for visual purposes. Number of dispersal events between the seven regions inferred using Biogeographical Stochastic Mapping on the DEC model **(b)**, and **(c)** number of transitions between open and closed habitats inferred to have occurred within each of the seven areas, calculated through comparison of best fitting corHMM models and historical biogeographical estimates (line thickness indicates relative frequency). The colours of row and column labels in **(b)** and arrows in **(c)** correspond to those on inset map in **(a)**.

**Figure 5.**
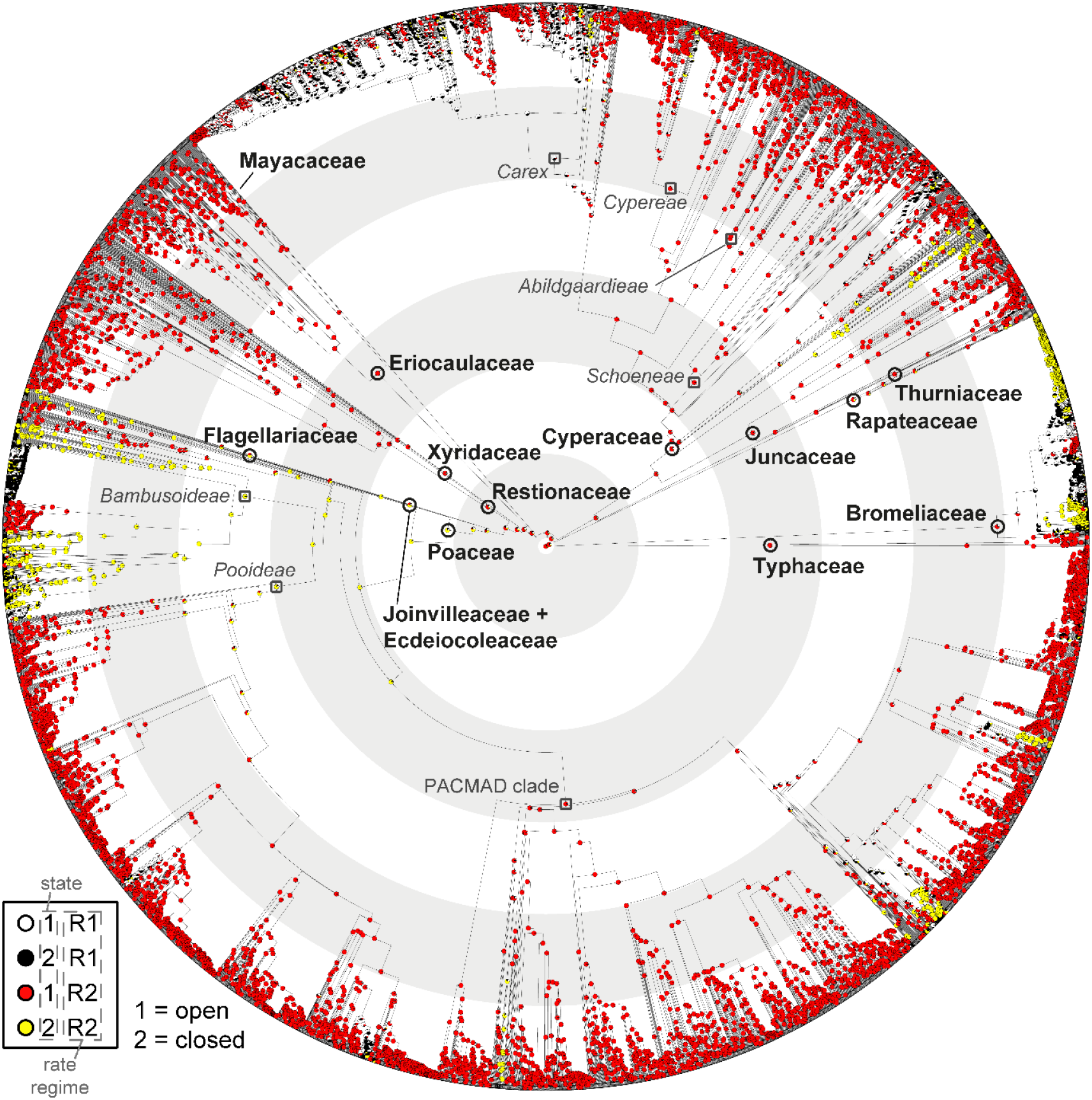
Ancestral states of open/closed habitats based on Generalized Hidden Markov models, with two rate regimes and asymmetric rates, implemented with the corHMM package in R. The crown nodes of the families within Poales are shown with black circles, whereas dark grey squares are used to depict lineages important for the study’s interpretations. Concentric light grey/white rings underlying the phylogeny indicate time slots of 20 million years intervals. Detailed transition rates between the states and rate are given in Table S5.

However, there are rapid rate transitions between open and closed (0.31 transitions / Mya in R1 [Regime rate 1]) and closed to open (0.51 transitions / Mya in R1) habitats among Bromeliaceae, Cyperaceae (*Carex* clade) and Poaceae (Bambusoideae clade), occurring predominantly during the Miocene.

## Discussion

This is the most comprehensive evolutionary study of Poales to date, with the goal of understanding the assembly of open habitats around the world. The key outcome is an emerging picture of parallel evolution – in space, through time, across lineages – resulting in the global assembly of open habitats, with notable, spatially and phylogenetically restricted divergences into strictly closed habitats. Our analyses show that each family exhibits a contrasting historical biogeography and pattern of habitat transition, which manifest in unique spatial patterns of assembly. And yet, patterns of evolution and assembly are repeated in most families.

### The biogeography and regionalization of Poales

The Poales represent over 120 million years of diversification, with an estimated Cretaceous origin in Western Gondwana (Fig. 4a). Consistent with previous phylogenetic studies, our model places the Poaceae origin in the Afrotropics with eventual migration to the Neotropics in the Eocene (Bouchenak-Khelladi *et al*., 2010; Gallaher *et al.,* 2022). Meanwhile Cyperaceae originates in the Neotropics and quickly shifts to Patagonia-Australia before landing in Africa in the Paleocene (Spalink *et al.,* 2016; Larridon *et al.,* 2021a). The earliest Cyperaceae fossils also place extinct Cyperaceae lineages in southeast Asia by the Paleogene (Smith *et al.,* 2009). The following global dominance of these Poaceae and Cyperaceae clades is a tale of parallel diversification with repeated spatial convergence of distantly related lineages, which is also observed in transitions between open and closed habitats and has observable impacts on spatial patterns of phylogenetic endemism.

In contrast to Poaceae and Cyperaceae, the other families have experienced more limited diversification, either spatially or in terms of species richness. The oldest of these is Restionaceae, which originated in Western Australia in the Late Cretaceous and dispersed to southern Africa in the Palaeocene (Fig. 4a; Linder *et al*,. 2003). The remarkable diversity of this family of C_3_ perennials remains restricted to open habitats in these two regions, where they have evolved strategies to withstand frequent fires (reseeder/resprouter; Litsios *et al*., 2014) and thrive on oligotrophic soils via enhanced nutrient acquisition strategies (cluster roots; Lambers *et al*., 2006), traits only shared by the Schoeneae (Cyperaceae; Barrett, 2013) and lacking in the Poaceae. Typhaceae and Juncaceae, both of Palearctic origin near the KP boundary, are now cosmopolitan but remain relatively species poor. Eriocaulaceae originated in the Neotropics at about the same time, and only a few lineages in the family have diversified outside of this ancestral zone. Bromeliaceae are perhaps the most unique among the Poales, diversifying nearly exclusively in Central and South America and with suites of traits not found in other families, such as epiphytism and lithophytism, which are uncommon traits in the Poales encountered outside the Bromeliaceae among the poikilohydric clade of Cyperaceae (Trilepideae; Muasya *et al*., 2010; Porembski *et al*., 2021).

These historical processes have resulted in substantial spatial structure to phylogenetic regionalization in Poales, with lineages forming thirteen distinct phyloregions (Fig. 2a) clustered into three floristic kingdoms (*sensu* Carta *et al*., 2022; SI Fig. S1). Despite the propensity for long distance dispersal in many Poales groups (Linder *et al*., 2018; Martín-Bravo *et al*., 2019; Benítez-Benítez *et al*., 2021; Larridon *et a*l., 2021a; Gallaher *et al*., 2022), regionalization in this clade largely mirrors that of all vascular plants, where the tectonic biogeographic legacy of Gondwana and Laurasia persists in determining, at least in part, the distribution of lineages (Takhtajan *et al*., 1986; Carta *et al*., 2022). In Poales, this is evident in northern temperate regions being distinct from those in the tropical south, and a united Austral-Patagonian (holantarctic, *sensu* Takhtajan *et al*., 1986) region. Unlike other vascular plants, temperate zones in both hemispheres share more common lineages than they do with tropical zones, and the Neotropics are phylogenetically distinct from the Paleotropics (SI Fig. S1). The unified North-South Temperate floristic kingdom may be partly due to bipolar disjunctions within species (at least 14) and genera (at least eight) in Poales (Villaverde *et al*., 2017). Likely, a more important driver of this pattern is the ecological filtering of lineages with respect to traits that evolved earlier in the diversification of Poales. These would include traits like frost hardiness and seasonality tolerance in Juncaceae, Typhaceae, and select Poaceae and Cyperaceae clades (Vigeland *et al*., 2013; Ambroise *et al.,* 2019; Martín-Bravo *et al*., 2019; Schubert *et al*., 2020), and edaphic specificity coupled with functional traits associated with adaptations to temperature and water extremes, as well as disturbance regimes (e.g., C_4_, CAM, silica, tannins, epiphytism; Givnish *et al*., 2011, 2014; Linder *et al*., 2018) across Poales. The relatively few lineages that have evolved tolerance to frost and strong climatic seasonality have diversified extensively, virtually in all habitable places where these conditions persist (Fig. 2). The phylogenetic differentiation of the Neo- and Palaeo- tropical kingdoms has roots deep in the Poales phylogeny. The Neotropical origin of Cyperaceae and Afrotropical origin of Poaceae, the near restriction of Eriocaulaceae, Bromeliaceae, and Rapateaceae in the Neotropics, and the restriction of Restionaceae to austral regions (Australia, New Zealand, southern Africa and Patagonia) drive strong phylogenetic beta diversity across the tropics.

### Evolutionary parallelism in open and closed habitats

The remarkable parallel evolution of open and closed lineages in Poales is most evident in Poaceae and Cyperaceae, which collectively represent ∼74% of species richness in the order and the majority of open-habitat PD (SI Fig. S3). Both families encompass two species-rich lineages (Cyperaceae: *Carex* and Cypereae; Poaceae: PACMAD and Pooideae), each of which (except Pooideae) is inferred to have an open-habitat ancestor (Fig. 5). The exclusively C_3_ *Carex* and Pooideae are predominately temperate lineages, likely originating in eastern Asia (Fig. 4; Gallaher *et al.,* 2022; Spalink *et al.,* 2016; Martín-Bravo *et al.,* 2019), while tropical Africa is inferred as the origin for the heavily C_4_ PACMAD and Cypereae clades (Figs. 4a, 5). These species-rich lineages evolved from C_3_ ancestors, acquiring cold tolerance (*Carex*, Pooideae; Vigeland *et al.,* 2013; Martín-Bravo *et al*., 2019; Schubert *et al*., 2020) and C_4_ photosynthesis (Cypereae, Abildgaardieae, PACMAD; Fig. 5) in parallel. The tempo of habitat transition has shifted from slow to fast multiple times (Fig. 5), most notably in *Carex* (> 2000 species; Larridon *et al.,* 2021b), Bambusoideae (c. 1700 species; Soreng *et al.,* 2022), and Bromeliaceae (c. 3700 species, Gouda & Butcher, 2023). Bambusoideae uniquely grow tall enough to reach the canopy (Soreng *et al.,* 2015; Attigala *et al*., 2016).

Bromeliaceae likely originated in open habitats, but has rapidly and repeatedly shifted between both habitats (Fig. 5).

Ultimately, our analyses indicate that there was likely a high diversity in all phyloregional species pools from multiple families with beneficial adaptations for survival in open habitats prior to their expansion in the Miocene (Edwards *et al*., 2010; Veldman *et al*., 2015). Consistent with the fossil record (Prasad e*t al*., 2005; Vicentini et al., 2008), the origin of PACMAD (encompassing all known origins of C_4_ in Poaceae; Edwards, 2019) and Abildgaardieae and Cypereae (majority of C_4_ Cyperaceae) is placed in the Palaeocene and Eocene (Fig. 4; Prasad *et al.,* 2011; Gallaher *et al.,* 2022), but most of the diversification in these clades coincides and is possibly linked with the low CO_2_ concentrations, cooler, drier conditions, and increased fire activity and herbivory during the Miocene (Osborne & Freckleton, 2009; Edwards & Smith, 2010; Strömberg, 2011; Maurin *et al*., 2014; Sage *et al*., 2018; Peppe *et al*., 2023). Diversification of most major Poales families increased in all phyloregions during this time, with open-habitat species accumulating most rapidly in the Miocene (Fig. 2).

### Spatial phylogenetics of Poales

The spatial structure of Poales phylogenetic diversity at the resolution presented here (Fig. 3, SI Fig. S3–S4) is ultimately the result of the biogeographical and ecological processes rather than local contemporary ecological processes (Meynard *et al*., 2013; Ross *et al*., 2021). The resulting diversity patterns are complex (Fig. 3). Each family exhibits peak PD (Fig. 2) in a different botanical country – usually, but not always in their phyloregion of origin (Fig. 4a). The PD of the order as a whole, and each family independently, defies a traditional latitudinal diversity gradient (Kreft & Jetz, 2007). Though this is the dominant trend in open habitat lineages, the PD of closed habitat lineages peaks in equatorial forests and decreases poleward (Fig. 3a,b). These patterns reflect the tremendous diversification of Poales in open habitats around the world (Figs. 2, 4, SI Fig. S3), with much more limited diversification of closed habitat lineages outside of ancestral ranges.

Patterns of diversity in Poales generally align with our stated hypotheses. PD peaks in the tropics, where most families originated. Hotspots of PE (Fig. 3c-e) are almost strictly a southern hemispheric phenomenon, and with the exception of California in North America, never occur north of 33°N at this spatial resolution. This pattern is likely driven both by the presence of spatially restricted families in the Southern Hemisphere (e.g., Bromeliaceae, Restionaceae, and Rapateaceae; Fig. 2b) and the propensity of lineages to migrate across the Holarctic (Donoghue & Smith, 2004).

To our knowledge, this is the first study coupling CANAPE with historical biogeographical estimations on a global scale. We find contrasting patterns of palaeo- and neo-endemism clearly reflect historical biogeographical processes at the family level (Fig. 3e). In Bromeliaceae, centres of palaeoendemism occur in the Guayana Shield – the site of putative origin of the family – whereas centres of neoendemism are restricted to the Brazilian Shield, where Bromelioideae subsequently radiated (Givnish *et al*., 2011). Eriocaulaceae mirror this pattern (Andrino *et al*., 2023), with the added area of super-endemism in India, where nearly 20% of the family occurs in the Western Ghats region alone (Sunil *et al*., 2015). In Restionaceae, Western Australia, the ancestral home of the family, hosts palaeoendemics, while neoendemics are restricted to South Africa (Linder *et al*., 2003). And finally, Poaceae and Cyperaceae show high levels of both endemism types around the world, reflecting their rapid migration early in their evolutionary history (Fig. 4) and *in situ* diversification across phyloregions (Fig. 2). In both families, tropical savannas and forested regions primarily harbour palaeoendemics, whereas more recent diversification is found frequently on islands – both young and old – and mountains. Significant palaeoendemism is observed within old landscapes with buffered climates (South America, southern Africa, India, and Australia; Hopper *et al*., 2016), while significant neoendemism is predominantly within regions that have experienced Miocene geomorphological evolution – especially in Andean (Gregory-Wodzicki, 2000) and Himalayan mountains (Wang *et al.,* 2022). Areas containing a mixture of both neo- and palaeoendemism are observed in regions where old and young landscapes are interspersed (Hopper *et al*., 2016; Barros-Souza & Borges, 2022; Vasconcelos *et al.,* 2020).

## Conclusions

The biogeography of individual families in the Poales is distinct from early in their evolution, such that few major clades originating in approximate synchrony do so in the same places.

This is particularly evident in Poaceae and Cyperaceae, whose evolution is like two dancers on a global stage - a biogeo(choreo)graphy - characterised by parallel movement through a shared temporality but with varying tempos, spatial staging, and ecological rhythms, ultimately resulting in a final tableau of global dominance. Both families originate in the late Cretaceous, but on either side of the widening Atlantic Ocean, dispersing and evolving in parallel to achieve cosmopolitan distribution. In contrast, the other families have experienced more limited, spatially restricted diversification. We identify latitudinal and longitudinal patterns of diversification, most evident in the phylogenetic distinctiveness of the temperate regions and also in the neotropics separating from the palaeotropics, with highest endemism found south of the Tropic of Cancer. Parallel evolution is observed in habitat transition, whose tempo has shifted from slow to fast multiple times especially since the Eocene, driven by repeated evolution of traits enabling colonisation of open and closed habitats in different latitudes.

## Data availability

The datasets analysed in this study are available as Supporting Information. The new Angiosperms353 raw reads are deposited in the European Nucleotide Archive (PRJEB35285).

## Supporting information

Supplementary figures and tables merged

## Acknowledgements

The authors acknowledge the team who delivered the WCVP on which many of the analyses performed in this study are based. This work was partly funded by grants from the Calleva Foundation to the Plant and Fungal Trees of Life Project (PAFTOL) at the Royal Botanic Gardens, Kew. The authors are thankful for the Research/Scientific Computing teams at The James Hutton Institute and NIAB for providing computational resources used for the phylogenomic analyses performed in the “UK’s Crop Diversity Bioinformatics HPC” (BBSRC grant BB/S019669/1). Computational resources were also supplied by the project ‘e-Infrastruktura CZ’ (e-INFRA CZ ID:90140) supported by the Ministry of Education, Youth and Sports of the Czech Republic. Lizzy Wenk extracted data from AusTraits. This work was supported in part by USDA National Institute of Food and Agriculture (McIntire Stennis Project 1018692) and NSF #1902064 to DS; UNAM–DGAPA–PAPIIT (No. IA202319), “Investigación Científica Básica” CONACYT (No. 286249) to CGM; MICINN- AEI to ME (PID2021-122715NB-I00) and SM-B (PID2020-113897GB-I00). KR thanks DGAPA-UNAM for a postdoctoral grant (2022). ARZ and JH were each funded by a “Future Leader in Plant and Fungal Science” fellowship from the Royal Botanic Gardens, Kew.

## Author contributions

AMM, TLE, DS and IL conceived the study. AMM, TLE, ME, DS, IL, JH, RLB, SM-B, JIM-C, CGM, ACM, KJR-S, DAZ, COA, DC, MSV, KLW and DAS collected data, while TLE, DS, JH, ARZ, and ME conducted the analyses. AMM, TLE and DS wrote the manuscript with input from all co-authors. JIM-C, DS, ME and RLB created the figures.

## Supporting Information (brief legends)

Additional Supporting Information may be found online in the Supporting Information section at the end of the article.

**Figure S1.** Poales botanical regions grouped into three ‘floristic kingdoms’ based on phylogenetic beta diversity, indicated by different colours and numbers.

**Figure S2.** The number of species of Poales missing from the phylogenetic dataset compared to the number listed in the World Checklist of Vascular Plants (WCVP) as of 28 February 2022, mapped per botanical region.

**Figure S3.** Phylogenetic diversity (PD) of the six largest Poales families categorised into open and closed habitats.

**Figure S4.** Phylogenetic endemicity (PE) mapped per botanical region for Poales and eight families with the highest number of species in the dataset.

**Fig. S5.** Ancestral area reconstruction within Poales based on seven regions, obtained using BayAREA-like model in BioGeoBEARS.

**Table S1.** Sequences for the phylogenetic reconstruction.

**Table S2.** Calibrations used in the treePL (Smith & O’Meara, 2012) configuration file.

**Table S3.** Taxa included in this study, showing habitat scoring across the Poales.

**Table S4.** Comparison of six ancestral area reconstruction models based on BioGeoBEARS analyses for Poales.

**Table S5.** Results from corHMM ancestral state reconstructions.

**Notes S1**. Justification for selecting dispersal-extinction-cladogeneis (DEC) model of ancestral estimation.

