## Supplementary figures and tables merged for "Global analysis of Poales diversification – parallel evolution in space and time into open and closed habitats"

### *1. Supporting Information: Figures*

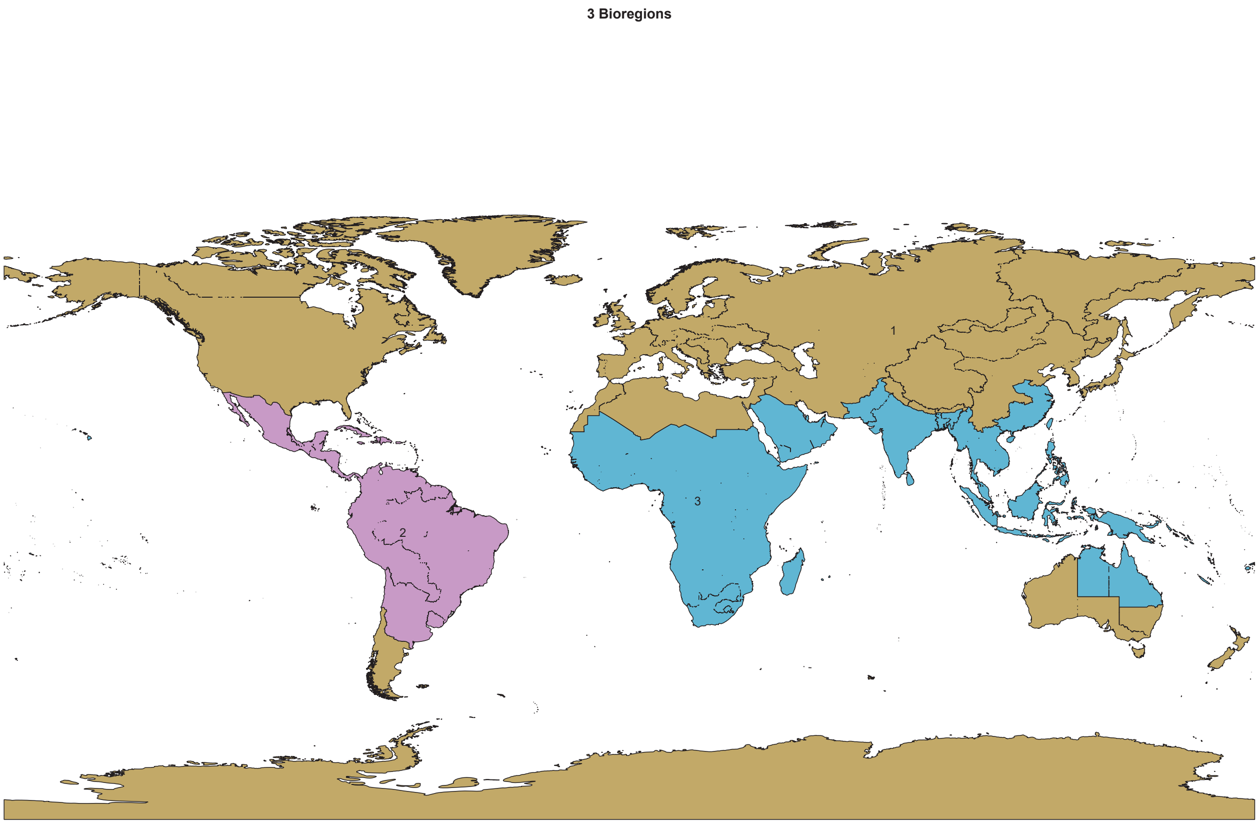


**Fig. S1** Poales botanical regions grouped into three 'floristic kingdoms' based on phylogenetic beta diversity, indicated by different colours and numbers.

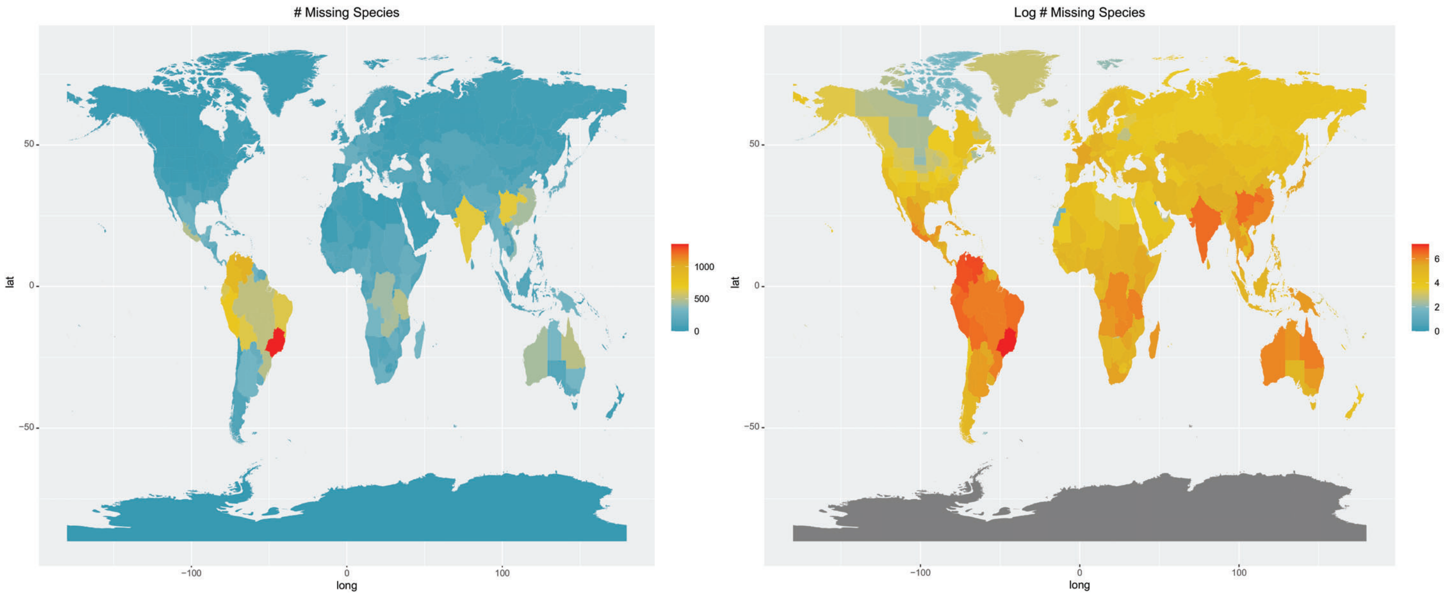


**Fig. S2** The number of species of Poales missing from the phylogenetic dataset compared to the number listed in the World Checklist of Vascular Plants (WCVP) as of 28 February 2022, mapped per botanical region. Red values indicate higher numbers of missing species, whereas blues represent less species missing from the dataset. The map on the left is based on untransformed values, while log-transformed values are mapped on the right hand side. The abbreviations ‘lat’ and ‘long’ denote latitude and longitude, respectively.

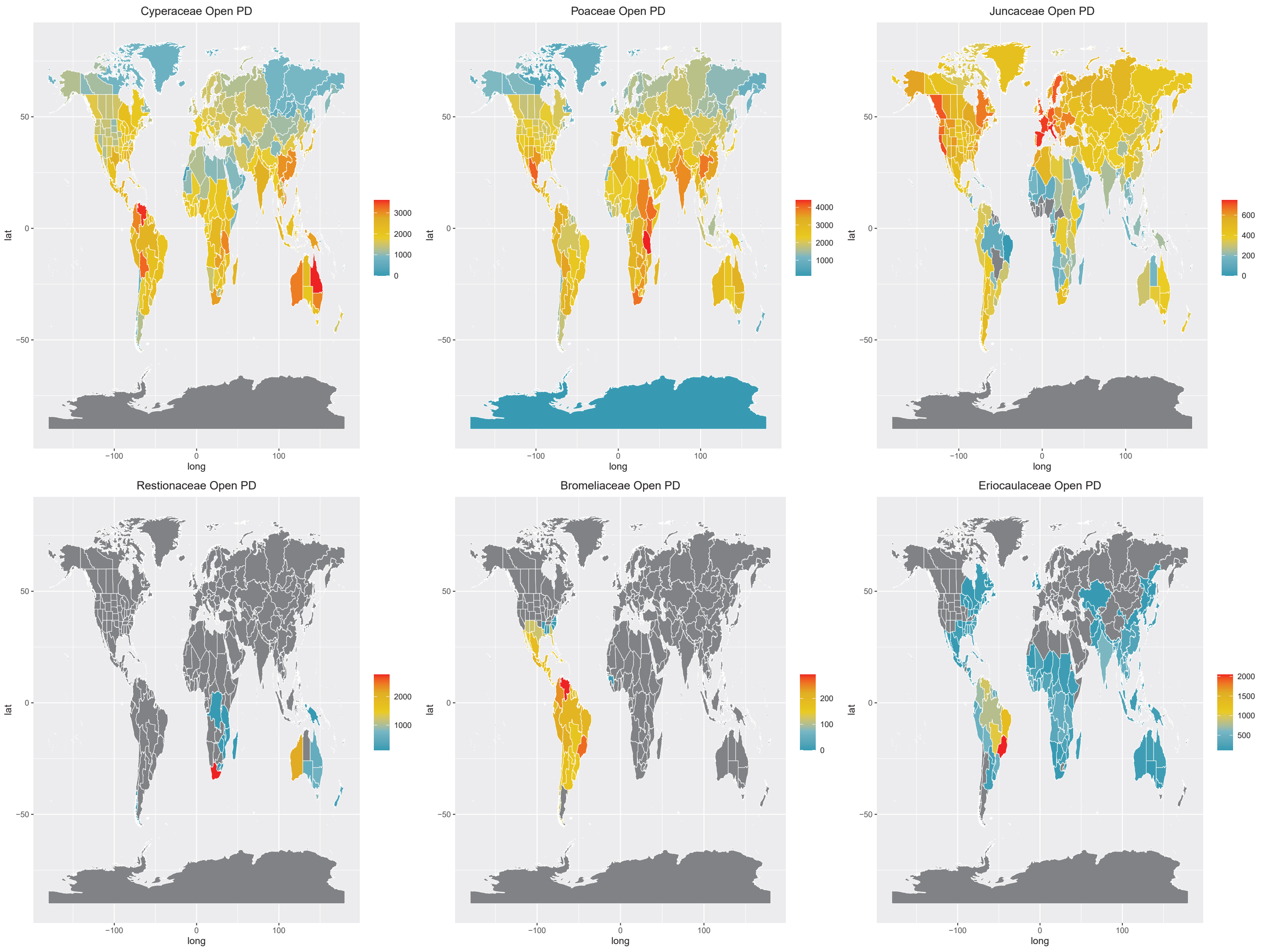


**Fig. S3** Phylogenetic diversity (PD) of the six largest Poales families categorised into open and closed habitats. Regions with high PD values for a trait are indicated in red, whereas low values are represented by blue values. Botanical regions without species for each clade are indicated by dark grey. The abbreviations ‘lat’ and ‘long’ denote latitude and longitude, respectively.

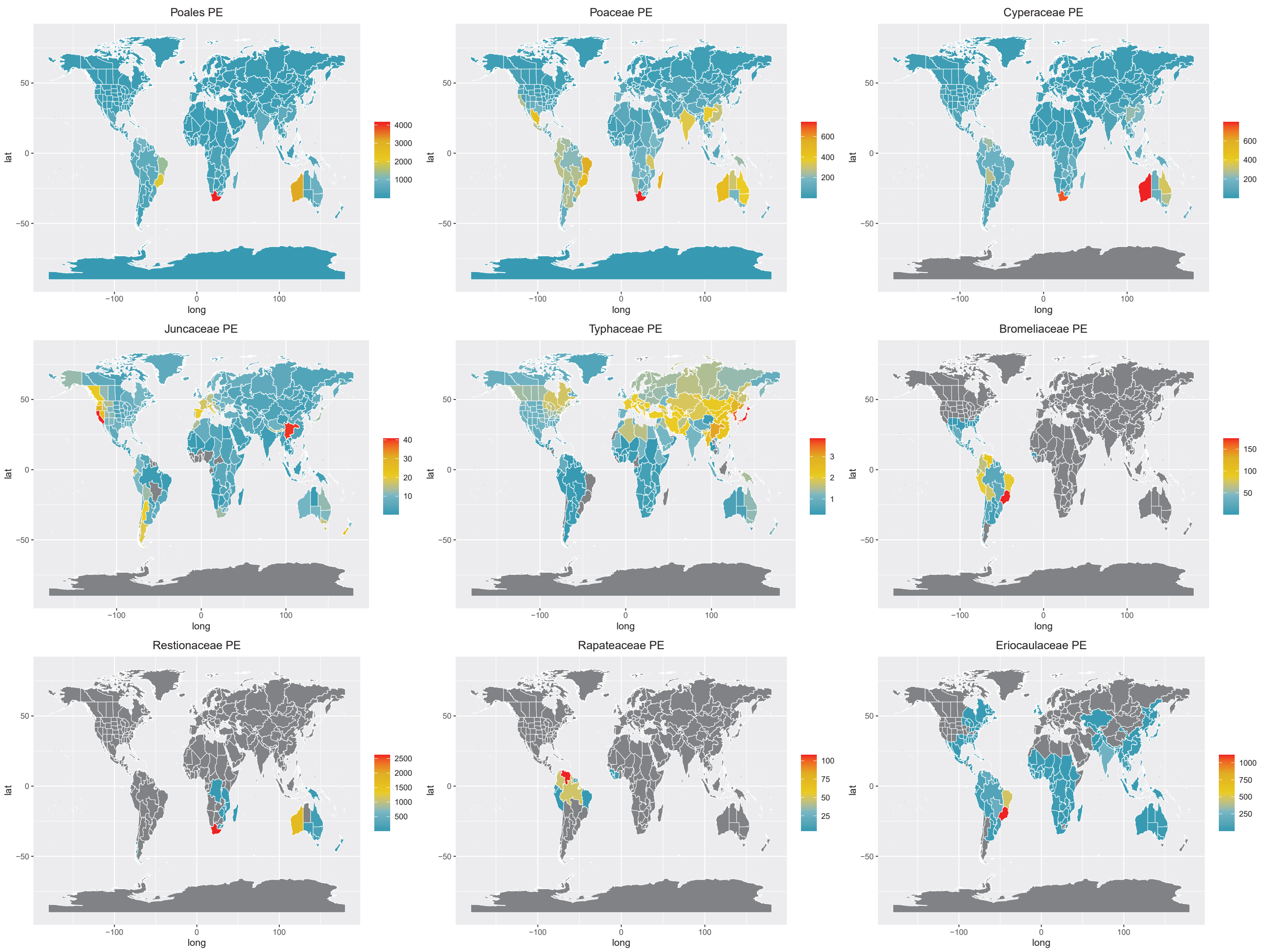


**Fig. S4** Phylogenetic endemicity (PE) mapped per botanical region for Poales and eight families with the highest number of species in the dataset. High PE values are indicated in red, whereas low values are represented by blue values. Botanical regions without species for each clade are indicated by dark grey. The abbreviations ‘lat’ and ‘long’ denote latitude and longitude, respectively.

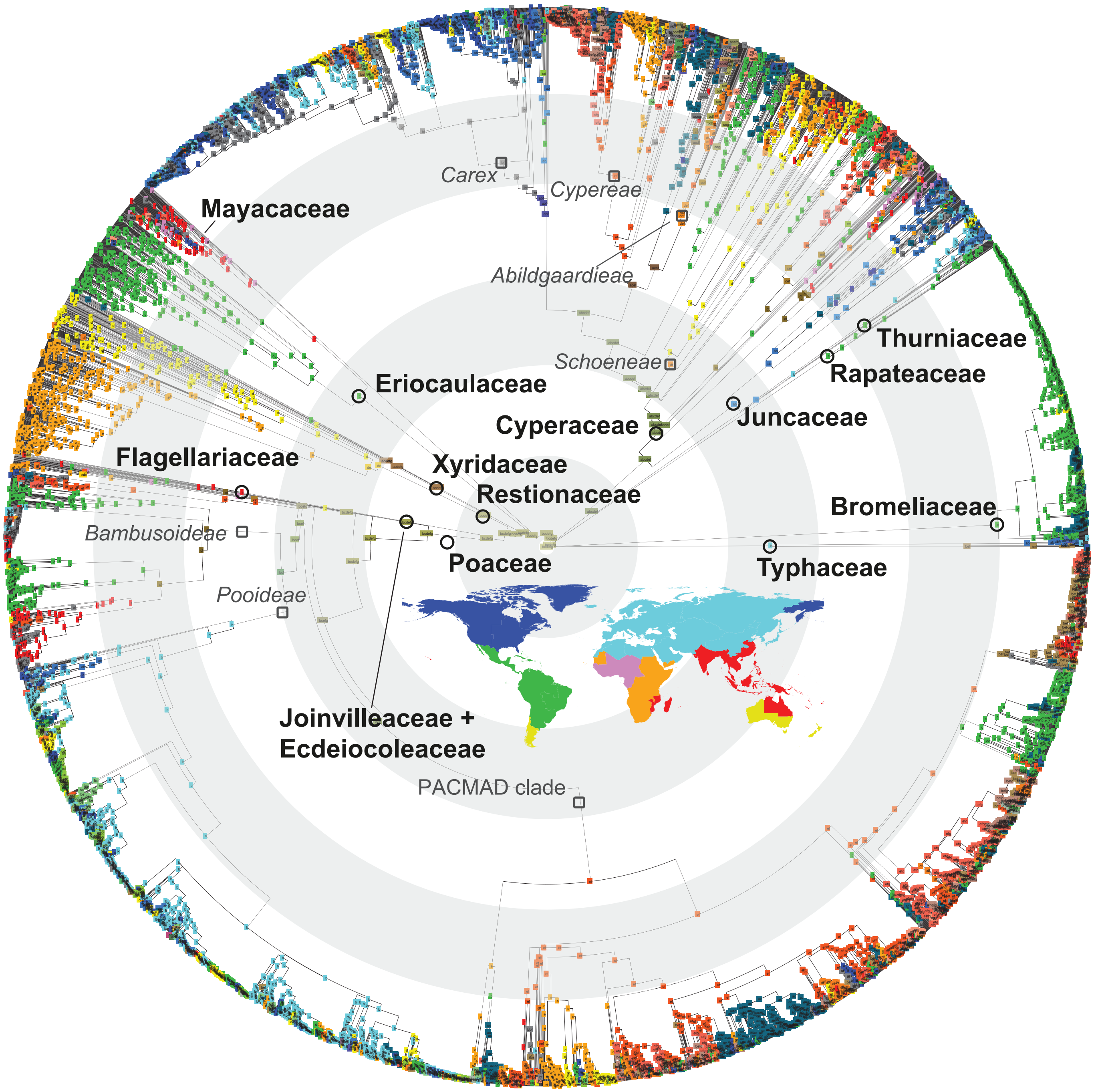


**Fig. S5** Ancestral area reconstruction within Poales based on seven regions, obtained using BayAREA-like model in BioGeoBEARS. Inset global map colours correspond to the seven defined areas for the BioGeoBEARS analysis. The crown nodes of the families within Poales are shown with black circles, whereas dark grey squares are used to depict lineages important for the study's interpretations. Concentric light grey/white rings underlying the phylogeny indicate time slots of 20 million years intervals. Note that Joinvilleaceae and Ecdeiocoleaceae are depicted together for visual purposes.

### 2. Supporting Information: Tables

**Table S2** Calibrations used in the treePL (Smith & O'Meara, 2012) configuration file.

| Clade | min | max | Backbone | tip_subclade1 | tip_subclade2 | Reference | Note |
|---|---|---|---|---|---|---|---|
| Poales | 120.1422 | 120.1422 | yes | (all) | (all) | Givnish *et al.* 2018 | Secondary; crown age used |
| Bromeliaceae | 19.9057 | 19.9057 | yes | Brocchinia_prismatica | Portea_fosteriana | Givnish *et al.* 2018 | Secondary; crown age used |
| Typhaceae | 70.3972 | 70.3972 | yes | *Typha* | *Sparganium* | Givnish *et al.* 2018 | Secondary; crown age used |
| Rapateaceae | 44.5638 | 44.5638 | yes | Stegolepis_hitchcockii | Rapatea_paludosa | Givnish *et al.* 2018 | Secondary; crown age used; *Potarophytum-Rapatea-Stegolepis* |
| Thurniaceae | 33.8687 | 33.8687 | no | Thurnia_sphaerocephala | Prionium_serratum | Givnish *et al.* 2018 | Secondary; crown age used; not constrained in backbone - only one tip |
| Juncaceae | 67.7516 | 67.7516 | yes | Juncus_pauciflorus | Luzula_elegans | Givnish *et al.* 2018 | Secondary; crown age used |
| Juncaceae_ Cyperaceae_ Thurniaceae | 107.3715 | 107.3715 | yes | Thurnia_sphaerocephala | Cyperus_diffusus | Givnish *et al.* 2018 | Secondary; crown age used |
| Juncaceae_ Cyperaceae_ | 90.8423 | 90.8423 | yes | Juncus_pauciflorus | Cyperus_diffusus | Givnish *et al.* 2018 | Secondary; crown age used |
| Xyridaceae | 92.5229 | 92.5229 | yes | Xyris_jupicai | Abolboda_grandis | Givnish *et al.* 2018 | Secondary; crown age used |
| Eriocaulaceae | 66.6136 | 66.6136 | yes | Comanthera_kegeliana | Eriocaulon_australe | Givnish *et al.* 2018 | Secondary; crown age used |
| Restionaceae | 104.6041 | 104.6041 | yes | Hopkinsia_anoectocolea | Restio_wittebergensis | Givnish *et al.* 2018 | Secondary; crown age used |
| graminids | 106.648 | 106.648 | yes | Flagellaria_neocaledonica | Andropogon_tracyi | Givnish *et al.* 2018 | Secondary; crown age used |
| Ecdeiocoleaceae | 70.1207 | 70.1207 | yes | Ecdeiocolea_monostachya | Georgeantha_hexandra | Givnish *et al.* 2018 | Secondary; crown age used |
| Cyperaceae | 75 | 88 | yes | Hypolytrum_longifolium | Schoenus_exilis | inclusive of Spalink *et al.*, 2016; Givnish *et al.*, 2018 | Secondary |
| *Carex* | 34 | 38 | no | Carex_moupinensis | Carex_longii | Jiménez-Mejías *et al.* (2016) | Fossil: *Carex colwellensis* |
| *Carex*_Vignea_clade | 16 | 23 | no | Carex_gibba | Carex_tribuloides | Jiménez-Mejías *et al.* (2016) | Fossil: *Carex marchica* |
| *Cyperus* | 24 | 32 | no | Cyperus_prolifer | Cyperus_nipponicus | Spalink *et al.*, 2016 | Secondary |
| *Eleocharis* | 31 | 41 | no | Eleocharis_robbinsii | Eleocharis_spiralis | Spalink *et al.*, 2016 | Secondary |
| *Fimbristylis* | 30 | 40 | no | Fimbristylis_compacta | Fimbristylis_densa | Spalink *et al.*, 2016 | Secondary |
| *Rhynchospora* | 40 | 50 | no | Rhynchospora_corniculata | Rhynchospora_megalocarpa | Spalink *et al.*, 2016 | Secondary |
| *Scleria* | 38 | 48 | no | Scleria_brownii | Scleria_virgata | Smith *et al.*, 2010 | Fossil |
| Poaceae | 75 | 95 | yes | Anomochloa_marantoidea | Andropogon_tracyi | Gallaher *et al.* 2022 | Secondary |
| BOP | 65 | 85 | no | Agrostis_lenis | Streptogyna_americana | Gallaher *et al.* 2022 | Secondary |
| PACMAD | 45 | 70 | no | Sartidia_jucunda | Andropogon_tracyi | Gallaher *et al.* 2022 | Secondary |

**Table S4** Comparison of six ancestral area reconstruction models based on BioGeoBEARS analyses for Poales. Models are listed based on ascending AICc values.

| Model | Log-likelihood | Number of parameters | Rate of dispersal | Rate of extinction | Jump dispersal at speciation | AICc |
|---|---|---|---|---|---|---|
| BAYAREALIKE + J | -20139 | 3 | 0.0079 | 0.026 | 0.0097 | 40284 |
| DEC | -21114 | 2 | 0.026 | 0.15 | 0 | 42233 |
| DEC + J | -21879 | 3 | 0.013 | 0.000000000001 | 0.0058 | 43765 |
| BAYAREALIKE | -22088 | 2 | 0.01 | 0.067 | 0 | 44180 |
| DIVALIKE + J | -23148 | 3 | 0.014 | 0.0000000033 | 0.007 | 46303 |
| DIVALIKE | -23831 | 2 | 0.017 | 0.0037 | 0 | 47666 |

AICc: corrected Akaike's information criterion; dispersal-extinction-cladogenesis (DEC); DIVALIKE (Dispersal-Vicariance Analysis); see Results section and **Notes S1** for justification of why DEC was chosen over BAYAREALIKE+J model.

**Table S5** Results from corHMM ancestral state reconstructions. Comparison between models **(a)** and parameters for the best fit model **(b)** are indicated.

**Table S5a** Comparison of corHMM models (2 rates vs. 1 rate and ARD vs. SYM) indicating AIC of the best model and deltaAIC in comparison with the best fit model.

| Trait | Model | AIC | deltaAIC |
|---|---|---|---|
| Open/Closed | 2 Rates ARD | 8814.2 | NA |
| | 1 Rate ARD | 9740.6 | 926.4 |
| | 2 Rates SYM | 9707.8 | 893.6 |
| | 1 Rate SYM | 10906.1 | 2091.9 |

**Table S5b** Model parameters associated with the 2 Rates ARD model.

| Trait | Model | 1R1 - 2R1 | 1R1 - 1R2 | 2R1 - 1R1 | 2R1 - 2R2 | 1R2 - 1R1 | 1 R2 - 2R2 | 2R2 - 2R1 | 2R2 - 1R2 |
|---|---|---|---|---|---|---|---|---|---|
| Open / Closed | 2 Rates ARD | 0.31 | 0.07 | 0.51 | 0.07 | 0.005 | 0.00000001 | 0.005 | 0.01 |

corHMM: Hidden Markov Models of Character Evolution; AIC: Akaike information criterion; ARD: all-rates-different; SYM: symmetrical model; 1: state 1; 2: state 2; R1: rate regime 1; R2; rate regime 2

The evolutionary history of important traits for Poales was reconstructed using Generalized Hidden Markov models, as implemented in the function corHMM of R package corHMM v.2.8 (Boyko & Beaulieu, 2021), to estimate the transition rates and ancestral state of several binary characters across the Poales tree phylogeny.

For each open / closed habitat binary trait, we ran the following Markov models:

- *symmetric rate* (SYM: one transition rate category; one parameter): transition rate (1 parameter); no hidden states
- *all rates differ* (ARD: one transition rate category; two parameters): transition rate for each regime (2 parameters); no hidden states
- *symmetric rate* (SYM: two transition rate categories; four parameters): transition rate for each regime (2 parameters); backward rate connecting two transition rate categories (1 parameter); forward rate connecting two transition rate categories (1 parameter); 2 hidden states; and
- *all rates differ* ARD: two transition rate categories; six parameters): two transition rates for each transition rate regime (4 parameters); two rates connecting the two transition rate categories (2 parameters); 2 hidden states.

**Notes S1** Justification for selecting dispersal-extinction-cladogeneis (DEC) model of ancestral estimation.

Because phyloregions are determined by spatial patterns of lineage turnover, they reflect – but do not necessarily conform to – discrete geologic boundaries typically used in ancestral area estimations (e.g., Martín-Bravo *et al*., 2019). However, these phyloregions present a data-driven hypothesis for the spatial relationship of areas as they relate to the biogeographical processes of dispersal and vicariance in Poales, and are thus well-suited for ancestral estimations. We a priori selected the dispersal-extinction-cladogeneis (DEC) model of ancestral estimation (Ree *et al*., 2005; Ree & Smith, 2008) instead of other available models (e.g., DIVA, Ronquist *et al*., 1997; BayArea, Landis *et al*., 2013), because our expectation is that the parameters of this model are best suited to the particular biology and distribution of Poales. For example, we expect both cladogenetic sympatry and vicariance to be important processes in Poales, particularly when descendent lineages diverge within only a portion of the ancestral range (i.e., subset sympatry) and when vicariant events unevenly split an ancestral range between two descendent ranges (i.e., narrow vicariance). The former scenario is not modelled by DIVA, while the latter is not modeled by BayArea. Many Poalean lineages are exceptionally good dispersers and able to migrate across typical migration barriers (e.g., oceans; Linder *et al*., 2018; Martín-Bravo *et al*., 2019, Spalink *et al*., 2019; Larridon *et al*., 2021, Benítez-Benítez *et al*., 2021), while lineages with species with poor dispersal ability tend to be restricted to single or physically adjacent phyloregions (e.g., Rapateaceae, Bromeliaceae). Highly parameterized models – with time-stratification or with geographic dispersal multipliers – are unlikely to be a good fit for all clades in the exceptionally diverse Poales.
